# PEGylated recombinant *Aplysia punctata* ink toxin depletes arginine and lysine and inhibits growth of tumor xenografts

**DOI:** 10.1101/2023.01.17.524398

**Authors:** Alena M. Wolkersdorfer, Birgit Bergmann, Juliane Adelmann, Matthias Ebbinghaus, Eckhard Günther, Marcus Gutmann, Lukas Hahn, Robert Hurwitz, Ralf Krähmer, Frank Leenders, Tessa Lühmann, Julia Schueler, Luisa Schmidt, Michael Teifel, Lorenz Meinel, Thomas Rudel

## Abstract

In recent years, a novel treatment method for cancer emerged, which is based on the starvation of tumors of amino acids like arginine. The deprivation of arginine in serum is based on enzymatic degradation and can be realized by arginine deaminases like the L-amino acid oxidase found in the ink toxin of the sea hare *Aplysia punctata*. Previously isolated from the ink, the L-amino acids oxidase was described to oxidate the essential amino acid L-lysine and L-arginine to their corresponding deaminated alpha-keto acids. Here, we present the recombinant production and functionalization of amino acid oxidase *Aplysia Punctata* ink toxin (APIT). PEGylated APIT (APIT-PEG) increased the blood circulation time. APIT-PEG treatment of patient-derived xenografted mice shows a significant dose dependent reduction of tumor growth over time mediated by amino acid starvation of the tumor. Treatment of mice with APIT-PEG which lead to deprivation of arginine was well tolerated.

## Introduction

Increasing resistance of tumors towards chemotherapies can be observed with standard of care DNA-damaging agents, antimetabolites, mitotic inhibitors, nucleotide analogous or inhibitors of topoisomerase^1^. The cytotoxic effect of the drugs is primarily by the induction of apoptosis. In treated tumor cells mutations of the general apoptosis signaling pathways can be observed, which leads to broad resistance to drug treatment due to the lag of activation of the apoptotic pathway^2^. This constitutes the need for alternative anti-tumor therapy approaches to tackle these limitations of current anti-tumor treatments.

An increasingly recognized strategy is the enzymatic depletion of amino acids in serum to target metabolic deficient malignances. A prime example of amino acid depletion cancer therapy is the treatment of leukemias with asparaginase. Normal cells convert aspartate to asparagine; however, this pathway is insufficient in leukemic cells. Treatment with asparaginase cures more than 90% of pediatric acute lymphoblastic leukemia patients^3, 4^.

Another highly discussed therapeutic approach is the depletion of arginine in arginine auxotroph tumors. Auxotrophy to the semi-essential amino acid arginine occurs through epigenetic silencing of ***a****rgino**s**uccinate **s**ynthetase 1* (ASS1) or ***a****rginine **l**yase (AL)* genes, preventing the synthesis of arginine through citrulline in the urea cycle^5, 6^. In many cancer types arginine-auxotrophy reaches an average of 60%-100%^7^. ASS1-deficient cells have a generally poor survival rate and an increased requirement for aspartate used for pyrimidine synthesis to support the proliferation of cells^8, 9^. This property can be therapeutically exploited by treating these tumor cells with **a**rginine **d**e**i**minases (ADI), which deaminate arginine to citrulline and thereby deplete arginine. In combination with arginine auxotrophy, cells are then starved from arginine and undergo apoptotic cell death^9, 10^. Therefore, a novel strategy for targeting auxotrophic cancers by starvation of the tumor of arginine emerged^11^.

A promising example for this treatment method is ADI-PEG20. First clinical trials with a mycoplasma-derived PEGylated ADI demonstrated the sensitivity of tumors towards arginine deprivation^12^. Based on this finding the FDA and the EMA granted orphan drug designation for ADI-PEG20 for the treatment of hepatocellular carcinoma and melanoma. Phase I clinical trials combined arginine deprivation with conventional cytostatically chemotherapy (cisplatin, pemetrexed-cisplatin, Nab-Paclitaxel with Gemcitabine, FOXLFOX6)^13^. In the studies patients showed an increase of immunogenic reaction against ADI-PEG20 after few weeks, which decreased the effectiveness of the combination therapy.

To overcome these limitations, the necessity of alternative products for ADI-PEG20 emerges including an arginine and lysine specific **L**-**a**mino **a**cid **o**xidases (LAAO) in the defense ink of *Aplysia punctata* (***A****plysia **p**unctata* **i**nk **t**oxin = APIT). APIT was first isolated from crude ink, which deaminates L-lysine and L-arginine to their corresponding α-keto acids^14^. APITs tumor specific cytotoxicity was demonstrated on apoptosis-resistant tumor cell^15^. However, in preliminary *in vivo* studies, intravenous injection of APIT into mice had an unfavorable renal clearance shortly after application^16^. To increase serum half-life, polymer modifications such as the coupling of **p**oly**e**thylene **g**lycol (PEG) to the protein are well-known strategies. Unspecific PEGylation of APIT, in which the lysine residues of the enzyme are targeted by N-hydroxysuccinimidyl chemistry, was performed, leading to increased serum half-life and exhibited comparable anticancer activity as ADI-PEG20^16^.

In this study, the effect of arginine and lysine deprivation mediated by APIT-PEG treatment was tested in the patient-derived HNXF 536 head and neck xenograft model (PDX). The results show that APIT-PEG efficiently and continuously depleted L-arginine and L-lysine levels in treated animals and significantly inhibited tumor growth in this model.

## Experimental Section

### Material and Methods

#### Recombinant expression of APIT

Competent *E. coli* HMS174 were transformed with the pET26b-based plasmid with inserted APIT encoding gene between *Nde*I and *Eco*RI restriction sites. Protein sequence of APIT is supplied in the SI. Expression was controlled by T7 promoter and induced by 1 mM Isopropyl-β-D-thiogalactopyranosid (IPTG) (Sigma Aldrich, Steinheim, Germany) at an OD600 of 0.8-1.0. After 4 h cells were harvested by centrifugation at 5000 g, 4°C. Expression resulted in inclusion bodies. Pelleted cells were resuspended in 50 mM Tris pH 8.0. Inclusion bodies were isolated in several sonification steps using a Bandelin Sonoplus ultrasonic device (Berlin, Germany).

#### Refolding of APIT

0.5 g of purified inclusion bodies were solubilized in 100 mL 6 M Urea, 50 mM **Tris**(hydroxymethyl)aminomethane (Tris), 200 mM NaCl, 100 mM Dithiothreitol (DTT) at pH 8.0 overnight at 4°C. Solubilized inclusion bodies were centrifuged at 5000 g, 4°C for 30 min and filtrated using a 0.2 µm Polyethersulfon (PS)-filter. The protein was diluted in refolding buffer 1 M Tris, 1 M NaCl, 1 mM EDTA, 25.6 µM FAD, 20 µM Cysteine, 0.5% PEG6000 pH 8.6 and stirred at 4°C for 7 days.

#### Purification of APIT

The subsequent purification of refolded APIT was performed in two steps using an anion exchange chromatography (HiTrap™ Q FF) and a size exclusion chromatography (HiLoad™ 16/600 Superdex™ 200 pg) from Cytiva (Uppsala, Sweden).

Elution was performed with buffer A (50 mM Tris, pH 8.0) and buffer B (50 mM Tris, 1 M NaCl, pH 8.0), starting with a washing step of 25% B in 10 CV, followed by an elution gradient of 25-60 in 10 CV. APIT was then loaded onto a HiLoad™ 16/600 Superdex 200 pg. Column was then run using Phosphate-buffered saline (PBS) pH 7.4 at flowrate of 1 mL/min.

#### PEGylation and purification of APIT-PEG5kDa

Since tris(2-carboxyethyl) phosphine (TCEP) added prior to storage of APIT at -80°C (see results section) interfered with the PEGylation reaction, APIT was dialyzed in PBS pH 7.4 overnight at 4°C and concentration was adjusted to 1.1 mg/mL. For the PEGylation of APIT, the 5 kDa linear form was selected in a preliminary experiment, in which the serum stability of native APIT modified with a variety of branched and different sized PEG forms was tested (Fig. S1). 20 eq of PEG-N-hydroxysuchinimid (NHS) 5kDa (Celares GmbH, Berlin, Germany) was dissolved at a concentration of 408 mg/mL in PBS pH 7.4 and 1:10 (v:v) added to APIT. The reaction was performed overnight at 4°C continuously shaking. Reaction was purified using an anion exchange chromatography (HiTrap™ Q HP, Cytiva, Uppsala, Sweden) with buffer A (50 mM Tris, pH 8.0) and buffer B (50 mM Tris, 1 M NaCl, pH 8.0). The reaction was diluted 1:3 in buffer A, a maximum of 10 mg of protein was loaded onto 1 mL of resin at a flowrate of 0.5 mL/min. Afterwards a washing step at 0% B for at least 10 CV and elution gradient from 0-100% B was performed. Elution peak was pooled and dialyzed against PBS pH 7.4, concentration was adjusted to 2 mg/mL PEGylated APIT was filtered using a 0.2 µm PS-filter and stored light protected.

#### Enzyme activity

The enzyme activity in the presence of the substate L-Lysine was determined by measuring the production of hydrogen peroxide. H2O2 is reduced stoichiometrically by the chromogenic 2,2’-Azino-di(3-ethylbenzthiazolin-6-sulfonic acid (ABTS) in a coupled Horseradish peroxidase (HRP)-catalyzed reaction as previously described by Butzke et al.^15^. The enzyme activity is expressed in units [U], 1 U is equivalent to the production of 1 µmol H2O2 per minute.

#### HPLC Analysis

Purity of produced monomeric APIT and PEGylated APIT was assessed by High performance liquid chromatography (HPLC) analysis using an Agilent Technologies 1100 series system (Waldbronn, Germany). A ZORBAX 300SB-CN, 300 A, 5 µm, 4.6 x 250 nm was used. For elution was performed with Water in HPLC grade + 0.1% Trifluoroacetic acid (TFA) and increasing concentrations of Acetonitrile HPLC grade + 0.1 TFA.

#### Degree of PEGylation

The degree of PEGylation was assessed by determination of the mass increase measured by Matrix assisted laser desorption ionization (MALDI) mass spectra with an ultrafleXtreme mass spectrometer (Bruker Daltonics, Bremen, Germany) equipped with a 355 nm smartbeam-II™ laser. 20-50 µg of protein were desalted using Sep-Pak Vac C18 cartridges. The recommended manufacturers instruction was followed. Bound protein was eluted using 80% acetonitrile in ddH2O+ 0.1% TFA and subsequently lyophilized at -100°C, 0.001 mbar. For external calibration Bruker Protein II standard (Bruker Daltonics #8207234) was mixed with Sinapinic acid (SA) solution (saturated in a 1:2 mixture (v/v) of Acetonitrile and 0.1% TFA). Dried drops were measured on a stainless-steel target (MTP 384 target plate ground steel, Bruker Daltonics #8280784). The proteins were analyzed after double layer preparation. First a thin matrix layer of SA solution (saturated in Ethanol) was applied onto the stainless-steel target, second the sample solution (4 mg/mL in a 1:1 mixture of Acetonitrile and ddH2O) was mixed with SA solution (saturated in a 1:2 mixture (v/v) of Acetonitrile and 0.1% TFA) and dropped onto the first layer. Several thousand shots were accumulated for the spectrum.

#### Pharmacodynamics of APIT-PEG on plasma L-arginine and L-lysine levels

Depletion of Arginine and Lysine in blood serum was assessed after a single intravenous administration of 250 U/kg of APIT, 1000 U/kg, 250 U/kg APIT-PEG or 250 mL/kg PBS as control vehicle into 5 tumor-free mice each group. Blood was collected 1 h prior to injection and 0.25 h, 1 h, 3 h, 6 h, 24 h and 48 h after injection from each mouse. Samples of each timepoint were pooled.

#### Ethics statement

Mouse experiments were approved by the German Committee on the Ethics of Animal Experiments of the regional council (Permit Numbers: G-21/100 and G-20/163). This study was carried out in strict accordance with the recommendations in the Guide for the Care and Use of Laboratory Animals of the Society of Laboratory Animals (GV SOLAS) in an AAALAC accredited animal facility.

#### Dose-finding experiments and efficacy experiments of APIT-PEG in HNXF 536 PDX mice model

The maximum tolerated dose (MTD) of APIT-PEG was evaluated at three dose levels of 15 U/kg, 30 U/kg and 50 U/kg in 4 tumor-free mice per group. APIT-PEG was intravenously injected every other day for one week and the tolerability was assessed by daily monitoring the mice for symptoms and body weight development.

Tumor fragments of HNXF 536 of a serial passage in nude mice were placed in PBS containing 10% penicillin/streptomycin and cut into fragments of 3-4 mm diameter. Prior to tumor implantation, NMRI nu/nu (Crl:NMRI-*Foxn1^nu^*) mice were anesthetized by inhalation of isoflurane. Animals were monitored until the tumor implants reached an appropriate size of 50-250 mm^3^.

Mice bearing the HNXF 536 were randomly divided into control and treatment groups. The day of randomization was designated as Day 0 of the experiment. Mice were treated with 1) control vehicle 5 ml/kg/day every other day 2) APIT-PEG 15 U/kg/day every other day 3) APIT-PEG 40 U/kg/day every other day 4) Doxorubicin-HCl 6 mg/kg every 7 days (See Tab. SI 1). Injection was performed via the lateral tail vein. To control Arginine and Lysine depletion during the therapy, blood was collected 6 h, 24 h and 48 h after first treatment.

#### Analysis of amino acids in serum

For analysis of Arginine and Lysine levels in serum, blood was collected for by retrobulbar sinus puncture under isoflurane anesthesia. Plasma was stored in lithium heparin vials on ice. Vials were centrifuged at 2,000 x g for 5 min at 4°C. Separated plasma was transferred to a cooled tube on ice. 60 µL sample was diluted with 15 µL precipitation buffer and stored at 5°C for 60 min to precipitate proteins present. After centrifugation at 2 000 x g for 5 min at 4°C, the supernatants were diluted 1:1 with sample dilution buffer containing 200 nmol/ml Norleucin, which was used as internal standard. For quantitative and qualitative analysis of the amino acids Serin, Alanine, Lysine and Arginine at 570 nm the ARACUS Amino Acid Analyzer was used (MembraPure, Hennigsdorf, Germany).

#### H&E staining of dissected tumors

Tumors were dissected at day 42 after randomization. Tumor blocks were cut into 5 µm sections using a Leica RM2135 microtome (Leica Biosystems, Wetzlar, Germany). Slides were dried overnight at 40°C. Prior to staining slides were heated at 60°C for 30 min. Paraffin was removed by using xylene and rehydrated in Ethanol solutions and washed with distilled water. Sections were then stained with Hematoxylin for 8 min and washed in with water for 15 min. Prior to the Eosin staining for 2 min, the slide was dipped in 96% Ethanol. The slide was washed with Water and dehydrated in 100% Ethanol and Xylene and mounted using the Merck Neo-Mount solution.

#### Statistics

Relative tumor growth over time was compared to Day 0. Statistical analysis of the curves was performed by fitting a linear regression and a comparison of the mean beta-coefficient by ordinary One-way ANOVA in a multiple comparison to the control group. Statistical analysis on specific days was performed on the relative tumor growth comparing the treatments separately to the control using the unpaired, nonparametric Mann-Whitney-U test. A p value less than 0.05 was considered significant (\**p* < 0.05, \*\**p* < 0.01, \*\*\**p* < 0.001).

## Results

### Expression and purification of monomeric APIT

We have previously demonstrated that the arginine and lysine deaminase APIT, an enzyme found in the *Aplysia punctata* ink toxin, induces cell death in cancer cells, but not in normal cells. To develop APIT for potential future pharmaceutical application, the serum half-life of APIT had to be increased through PEGylation. We therefore developed the recombinant expression and subsequent purification of APIT, which resulted in approximately 8 mg/L monomeric APIT after the last purification step via size-exclusion chromatography. The most crucial part of the purification was the purity of the inclusion bodies prior to refolding, which was optimized by a serial of sonification steps. Refolded APIT was purified via anion exchange chromatography, which still contained high molecular APIT (Fig. 1A). To obtain pure monomeric APIT, a reductive step using tris(2-carboxyethyl) phosphine (TCEP) was introduced prior to size exclusion chromatography (SEC) (Fig. 1B). Over time, monomeric APIT tended to form dimers, which did not interfere with the enzymatic activity. To avoid dimerization, monomeric APIT was stored at -80°C with the addition of small amounts of TCEP. Monomeric APIT had a mass of approximately 59592.4 g/mol and a purity of ≥98% measured by HPLC (Fig. 1 C), with a small amount of dimeric APIT with a mass of 121457.8 g/mol (Fig. 1D) and a batch dependent enzyme activity of 80-100 U/mg.

**Figure 1.**
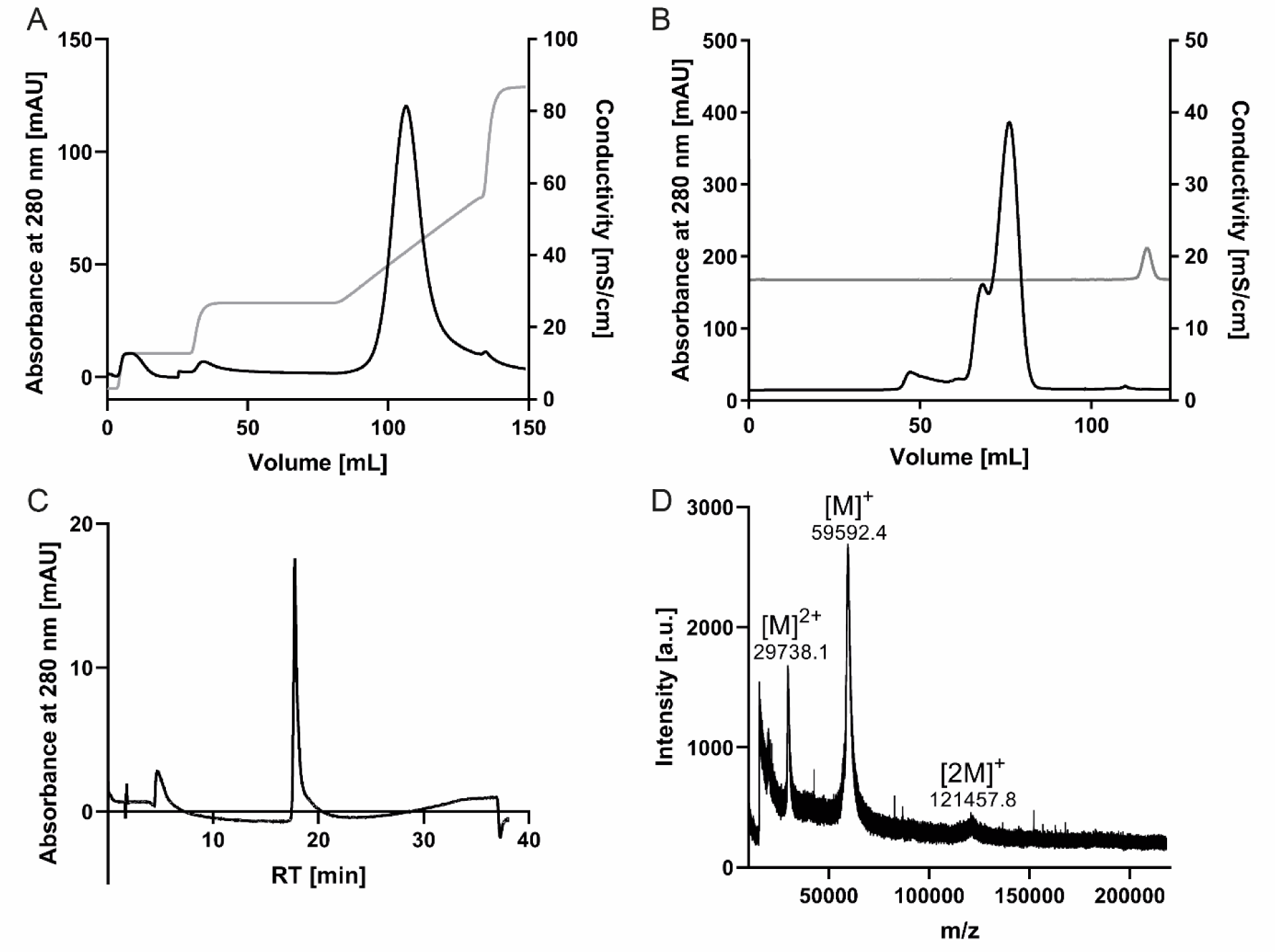
Purification and characterization of recombinant APIT A) Chromatogram of anion exchange after refolding of APIT B) Size-exclusion chromatogram of pooled and reduced APIT C) HPLC analysis of monomeric recombinant APIT after purification on a normal phase column D) MALDI-Mass of purified APIT showing small amounts of remaining dimeric APIT.

### Production of APIT-PEG for *in vivo* application

To enhance the serum half-life, we PEGylated purified APIT at lysine residues and purified the conjugates via anion-exchange chromatography (Fig. 2A). After conjugation, a protein recovery of 60% of pure APIT-PEG was achieved. Compared to APIT the enzyme activity of APIT-PEG was reduced by 20%. Purity of APIT-PEG was ≥95% measured by HPLC and SDS-PAGE (Fig. 2B-C). APIT-PEG showed an average increase of mass by 55 507 g/mol indicating a degree of PEGylation of approximately 8-10 mol PEG/ mol measured by MALDI-MS (Fig. 2D). APIT-PEG was formulated in PBS pH 7.2 and stored at 4°C. Prior to release for animal experiments, the endotoxin level was determined and shown to have a concentration of 9 EU/mg, 0.14 EU/U. Masking of endotoxin by formulated APIT-PEG was controlled by the spike-test. Spike recovery was 103-110%, concluding formulated APIT-PEG is not interacting with endotoxins and therefore hiding endotoxins.

**Figure 2.**
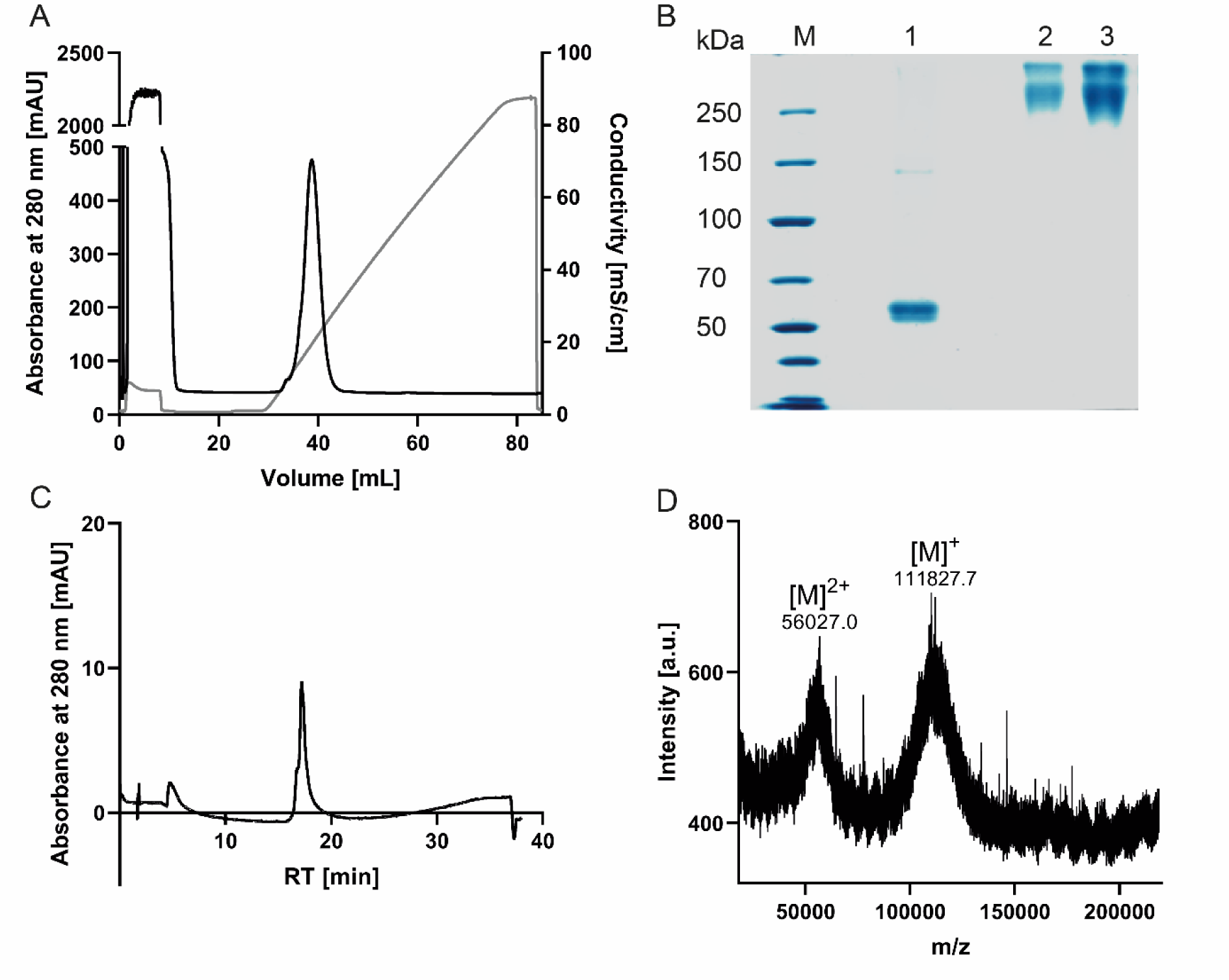
Purification of PEGylated APIT A) Chromatogram of Sepharose Q HP column B) 5-15% Tris-SDS-PAGE of non-reducing APIT (1) and APIT-PEG 2 µg (2) 4 µg (3) stained with Coomassie blue G250 C) CN-HPLC profile of PEGylated APIT after purification D) MALDI-mass spectrum of PEGylated APIT shows an increase of mass by 55507 g/mol, indicating a degree of PEGylation of 8-10 PEG per APIT.

### PEGylated APIT depletes serum arginine and lysine for at least 48 h

Depletion of arginine and lysine in mice blood was tested after a single injection of A) 250 U/kg APIT B) 250 U/kg APIT-PEG or C) 25 mL/kg PBS as control vehicle for dilution effects. Serum of mice treated with APIT showed a beginning recovery of arginine (Fig. 3 A) and lysine levels after 6 h (Fig. 3 B). A single injection of 1000 U/kg APIT showed the depletion of arginine and lysin for up to 6 h. Nevertheless, after 24 h a complete recovery of amino acids in serum were observable (Fig. S2). Only the injection of APIT-PEG showed a complete depletion of serum arginine and lysine during the 48 h period (Fig. 3).

**Figure 3.**
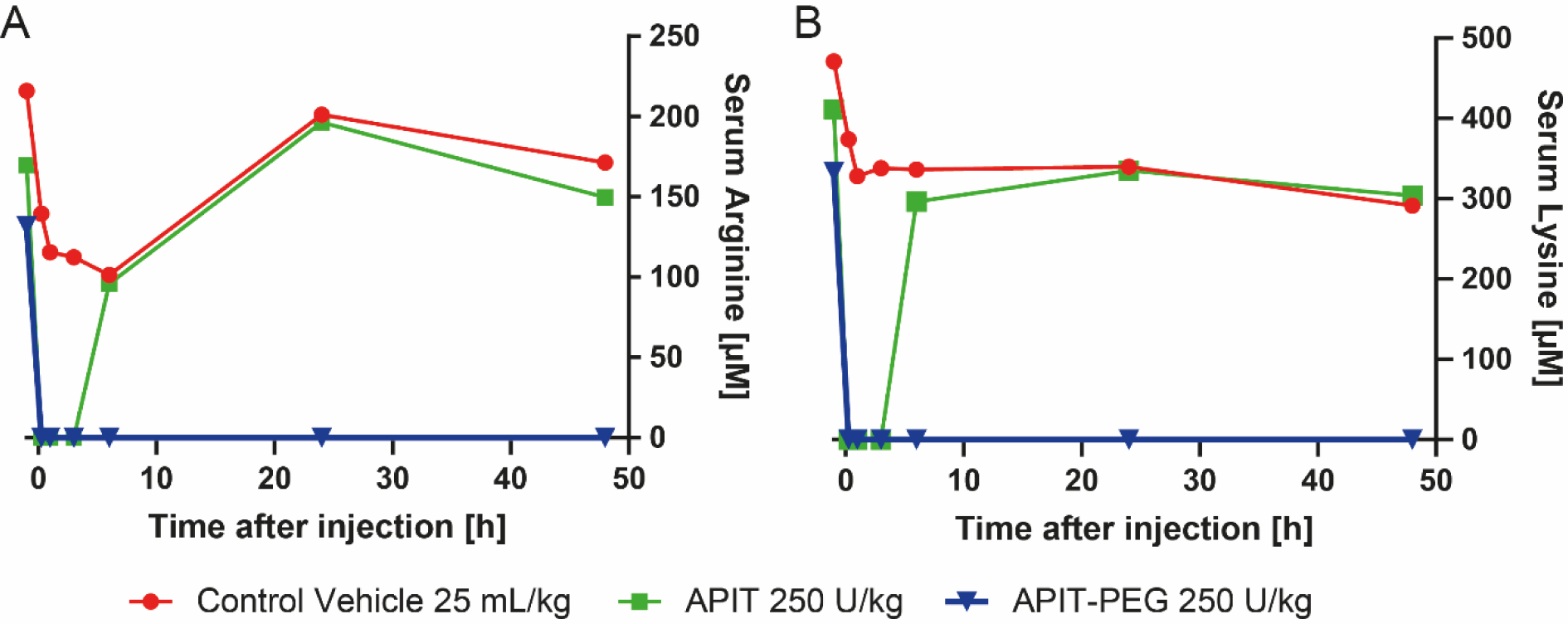
Pharmacodynamics of a single administration of 250 U/kg APIT or 250 U/kg APIT-PEG on A) arginine concentration in Serum and B) lysine concentration in serum showing beginning recovery of arginine and lysine concentration after 6 h, whereas APIT-PEG showed a complete depletion of serum arginine and lysine at least 48 h.

### Tolerability experiments in tumor-free nu/nu mice

Dose finding tolerability tests were performed to define the suitable dose that does not affect the animal’s health during the continuous treatment with APIT-PEG. Therefore, the relative change in body weight was taken as an indicator of tolerability and health since weight loss or gain should not exceed 10% in seven days during chemotherapy. Mice were treated with 15 U/kg, 30 U/kg or 50 U/kg of APIT-PEG every other day. Due to a high variance of body weight changes of the mice, especially in the group of the highest dose, the median of relative body weight change with a 5 to 95% confidence interval was used for evaluation to accurately depict results. The mice administered with 50 U/kg APIT-PEG showed a median body weight loss of 7.3% after one week, with one mouse above the limit of 10% showing a loss of 14.3%. The dose of 50 U/kg APIT-PEG was therefore declared as the upper limit (Fig. 4). The results of these experiments demonstrated that doses up to 50 U/kg were well tolerated.

**Figure 4.**
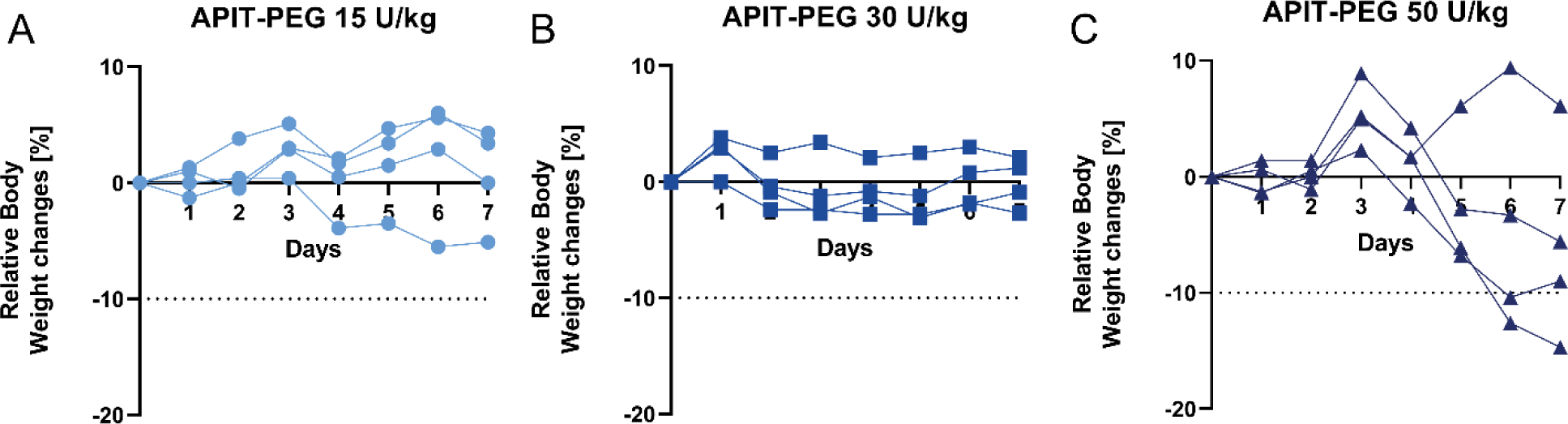
The median and 5-95% confidence interval (CI) of relative body weight change of mice treated with APIT-PEG every other day for one week shows a stronger impact of higher doses of APIT-PEG on the well-being of mice.

### Efficacy experiment of APIT-PEG in the HNXF 536 PDX mouse model

To evaluate the anti-tumor effect of arginine and lysine deprivation, we tested APIT-PEG in mice xenografted with HNXF 536, a patient-derived head and neck cancer (Fig. S3). Based on the tolerability experiment APIT-PEG was applied intravenously at 15 U/kg and 40 U/kg instead of 50 U/kg, since individual mice of the 50 U/kg group lost more weight than the limit of 10% (Fig. 4). APIT-PEG was administered every other day in this efficacy experiment in mice and doxorubicin as standard-of-care drug once a week at a concentration of 6 mg/kg. The control group received PBS every other day (for details see Tab. SI 1). At day 21 the treatment was terminated due to the high tumor load in the control group and lower dose group of 15 U/kg APIT-PEG. Tumor growth was monitored until day 28 and last observations were carried forward. Mice were weighed until day 42, relative body weight of doxorubicin showed a loss of weight during the treatment, the well-known impact of standard chemotherapeutic treatment approaches on health (Fig. 5A). Mice slowly regained body weight after the end of the treatment. APIT-PEG treatment in contrast did not result in severe changes of body weight and did not affect the mice’s health, when compared to the control group. After the end of the treatment all mice gained weight comparable to the control group. The tumor volume was determined from all mice as a measure for tumor growth (Fig. 5B). Treatment with doxorubicin resulted in significant loss of tumor volume starting at day 10. Treatment with 15 U/kg APIT-PEG did not show significant difference in relative tumor growth compared to the control group (Fig. 5B-D). Serum analysis of amino acids showed an increase of serum arginine after 48 h, probably leading to the lag of effectiveness due to incomplete arginine and lysine depletion (Fig. S4, 5). In contrast, treatment with 40 U/kg APIT-PEG successfully depleted arginine and lysine in blood serum during the treatment. Statistical analysis of the curves (Fig. 5B) was performed by fitting a linear regression and a comparison of the mean beta-coefficient by ordinary One-way ANOVA in a multiple comparison to the control group (Fig. S6). The curve of the relative tumor growth of 15 U/kg was not significant, whereas the 40 U/kg (p = 0.0493) and doxorubicin (p < 0.0001) treatment curves were significantly different to the control group. Evaluation of the tumor size on the last day of injection (*p*= 0.0160) and one week after the end of treatment (*p*= 0.0075) confirmed significant reduction in the 40 U/kg APIT-PEG group compared to the non-treated control group (Fig. 5C-D).

**Figure 5.**
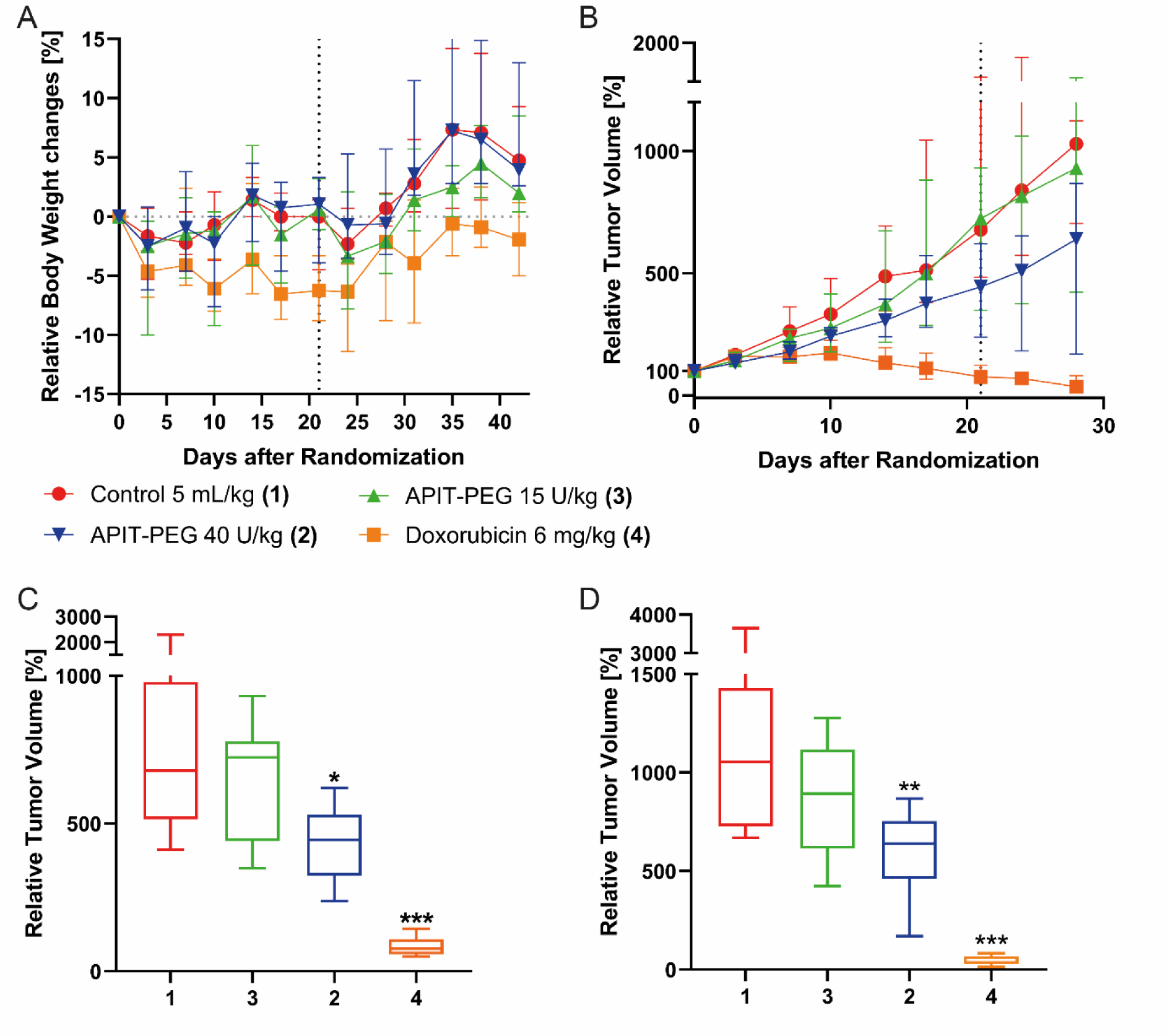
Efficacy experiment of treatment of patient derived HNXF 536 xenografted mice with APIT-PEG or Doxorubicin the end of treatments is indicated with dashes lines A) Median and 5-95% CI of relative body weight changes for monitoring the mice’s health B) Median and 5-95% CI of relative tumor growth during and one week after the treatment C) Boxplot representation of median and 5-95% confidence interval of relative tumor growth on day 21 (last day of treatment) D) and one week after the end of treatment on day 28. (Whiskers indicate 5-95% percentile values, the line in the middle of the box represents the median; Significance was tested with the Mann-Whitney U test, \**p* < 0.05, \*\**p* < 0.01, *** *p* < 0.001).

## Discussion

Targeting amino acid metabolism is an attractive anticancer strategy since it may provide alternatives for treatment of therapy resistant tumors (for review see^11^). Killing of therapy resistant tumor cell lines has also been demonstrated for APIT, which kills cancer cells independent of the induction of apoptosis^15^, the mechanism induced by many of the genotoxic agents used for cancer therapy^1^. Resistances of tumors to these canonical therapies occur frequently by mutations in the general apoptosis signaling pathways^2^. This explains why anti-tumor therapies provoke the emergence of broad resistance to drugs with the same mode of action, the induction of apoptotic cell death. In addition, investigations in amino acid depletion cancer therapies suggest that side effects like those elicited by current treatments with genotoxic agents are less severe or even not observed, since amino acid auxotrophies are often highly tumor specific^3^. A prime example of amino acid depletion cancer therapy is the treatment of leukemias with asparaginase curing more than 90% of pediatric acute lymphoblastic leukemia patients^4^.

Since APIT depletes the essential amino acid lysine and the semi-essential amino acid arginine^14^, both activities interfere with tumor cell amino acid metabolism. Besides increased need of proliferating cells for lysine, the enzymes involved in *de novo* synthesis of arginine, argininosuccinate synthetase (ASS1) and argininosuccinate lyase (ASL) are frequently deregulated in cancer cells^5^, leading to arginine auxotrophies in many different tumor cells (for review see^7^) This metabolic Achilles heel of tumor cells can be used for therapeutic approaches based on the activities of APIT. However, the blood circulation time of active purified APIT is insufficient to achieve the long-lasting depletion of L-lysine and L-arginine that is required to starve tumor cells.

Here we developed the PEGylation of purified APIT and thereby achieved a dramatic increase in the blood circulation time of the enzyme. A single dose of PEG-APIT depleted L-lysine and L-arginine for more than 50 hours whereas the amino acid oxidase activity of non-PEGylated APIT was cleared in less than 10 hours. Despite the long-lasting amino acid depletion, APIT-PEG treatment had no negative effect on the well-being of mice, whereas doxorubicin treatment resulted in weight loss. APIT-PEG induced a dose-dependent, significant decrease of the head and neck cancer HNXF 536 in patient derived xenografted mice compared to the control group. Interestingly, treatment with the lower dose of 15 U/kg APIT-PEG caused a non-significant reduction of the tumor volume and this correlated with a low but measurable level of L-arginine and L-lysine in the plasma of these mice. This is in line with the requirement of a long-lasting efficient depletion of L-lysine and L-arginine to achieve an anti-tumor effect. Due to the very good tolerability of the APIT-PEG treatment, the dose and thus the efficacy could be optimized in further experiments. Due to its broader action by depleting of two amino acids instead of only one, APIT-PEG could be an interesting alternative for ADI-PEG20, which was granted the orphan drug status and is currently used for the treatment of malignant pleural mesothelioma.

APIT was PEGylated on lysine residues using monofunctionalized NHS-activated polyethylene glycol. The high PEG concentration during the PEGylation resulted in minor precipitation of protein, explaining the protein recovery of 60% APIT-PEG. PEGylated APIT was successfully isolated by anion-exchange chromatography and contained between 8-10 mol PEG per mol APIT. Furthermore, the enzyme activity of APIT-PEG was about 20% reduced compared to APIT, indicating that PEGylation causes only a minor loss of enzyme activity.

For the current experiments, the purification yield was 8 mg monomeric APIT per liter culture with a batch-dependent enzyme activity between 80-100 U/mg. Further optimization of expression and large-scale production using fermenters will enable the production of PEG-APIT at a scale required for clinical testing. So far, a size-exclusion chromatography step is necessary to yield pure monomeric APIT. Further optimization to exclude higher molecular APIT might be achieved by selection of suitable hydrophobic interaction chromatography media to improve the purification procedure also for large scale approaches. Furthermore, a critical step for the application in clinical testing is the endotoxin amount. Reduction of endotoxin might be achieved by endotoxin reduced workflow via anion exchange chromatography and could be integrated in the existing purification steps in larger scale productions. Our work therefore gives us hope that preclinical testing will soon be completed and manufacturing and clinical testing can begin.

## Conclusions

This study presents the recombinant synthesis and purification of Arginine- and Lysine-deaminase *Aplysia Punctata* Ink Toxin (APIT). Due to a low serum half-life, PEGylation of lysine residues of APIT was performed. Increased serum half-life was proven in an *in vivo* study. Further, APIT-PEG was tested *in vivo* in a dose-escalation study to determine tolerated doses in healthy mice. Subsequently, the effect of arginine and lysine depletion in serum of mice carrying patient derived head and neck cancer resulted in a dose-dependent inhibition of tumor growth. In conclusion, this initial proof of principle study demonstrated the effect of APIT-PEG as a potential tumor therapeutic agent.

## Supporting Information Available

The following files are available free of charge:

Figure S1: Screening of APIT-PEG conjugates on the pharmacodynamic

Figure S2: Pharmacodynamics of administration of APIT and APIT-PEG on arginine and lysine concentration in serum.

Figure S3: Representative histology of patient derived head and neck cancer as xenograft in mice.

Figure S4: Amino acid concentration in serum of treated xenografted mice.

Figure S5: Serin, alanine, lysine, and arginine concentration in serum of different treated groups of xenografted mice.

Figure S6: Comparison of beta coefficients of the curves of the relative tumor growth performed by a linear regression fit.

## Acknowledgments

We thank Prof. Dr. Thomas Dandekar (Chair of Bioinformatics) for the help with the statistical analysis, and Prof. Dr. Andreas Rosenwald (Institute for Pathology) for the help with the analysis of histological sections.

## Authors’ contributions

Wolkersdorfer A.M.: Investigation, Writing original draft

Bergmann B.: Investigation

Adelmann J.: Validation, Resources

Ebbinghaus, M.: Investigation

Günther, E.: Funding Acquisition

Gutmann M.: Validation, Resources

Hahn L.: Validation, Resources

Hurwitz R.: Conceptualization, Methodology

Krähmer R.: PEGylation, Methodology

Leenders F.: PEGylation, Methodology

Lühmann T.: Supervision

Schueler J.: Investigation

Schmidt L.: Validation

Teifel M.: Project administration

Meinel L.: Supervision, Funding acquisition, Writing – Review & Editing

Rudel T.: Conceptualization, Project administration, Funding acquisition, Writing – Review & Editing

## Table of Contents graphic

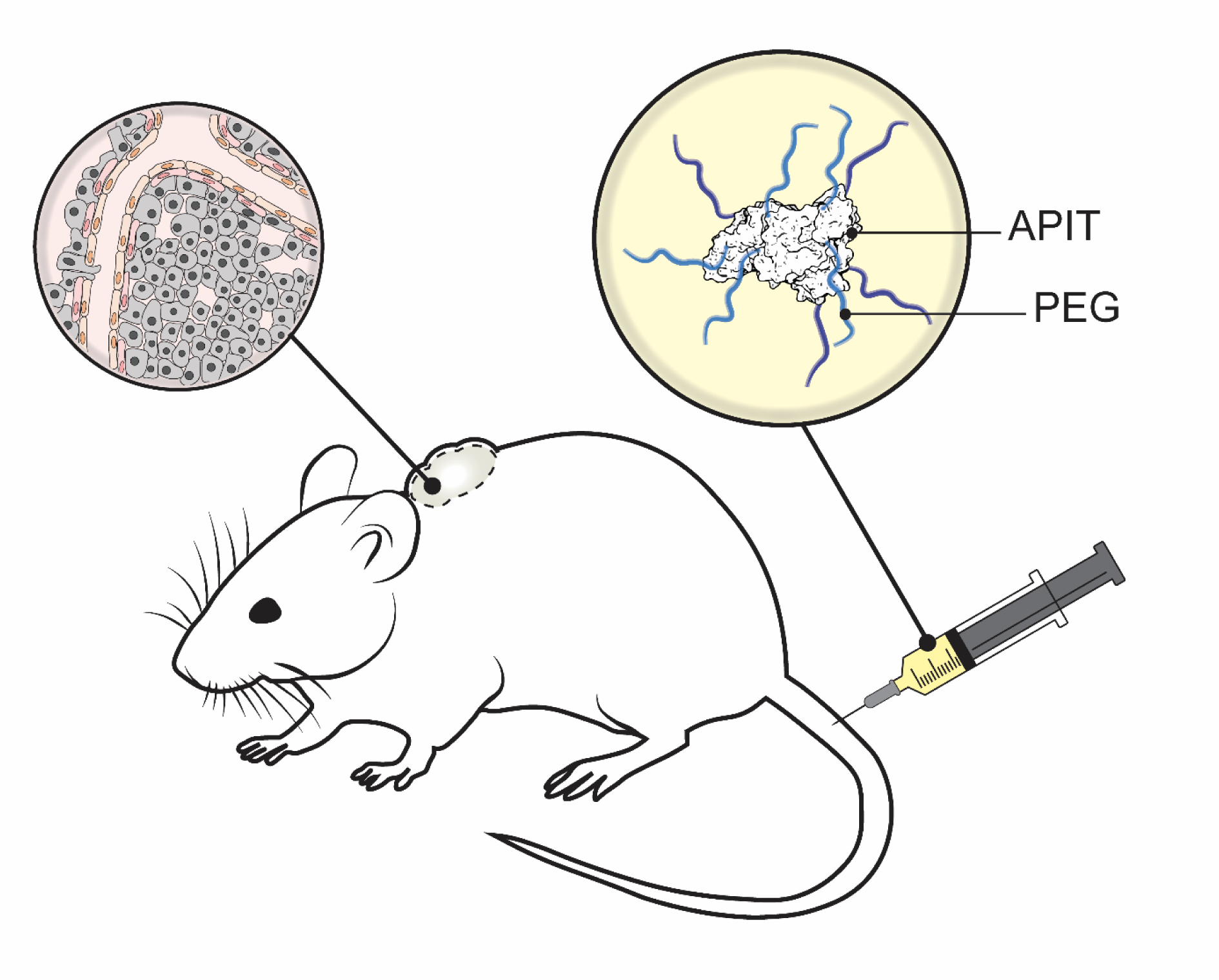

