## Supplemental Information for "PEGylated recombinant *Aplysia punctata* ink toxin depletes arginine and lysine and inhibits growth of tumor xenografts"

**Material**

Polyethylenglycol functionalized with N-hydroxysuccinimid with an average mass of 5 000 g/mol (PEG-NHS 5kDa) was purchased by Celares GmbH (Berlin,Germany). Isopropyl-ß-D-thiogalactopyranosid (IPTG), phosphate buffered saline (PBS), trifluoracetic acid (TFA, analytical grade), 2,2’-azinobis-(3-ethylbenzthiazolin-6-sulfonic acid)-tablets (ABTS-tablets, 10 mg) and horseradish peroxidase (HRP, Type VI-A for ABTS) were purchased by Sigma-Aldrich (Steinheim, Germany). EndoZyme® II kit was purchased of bioMerieux Deutschland GmbH (Bernried, Germany). Copper-based bicinchoninic acid (BCA)-Assay was purchased from ThermoFisher Scientific (Dreieich, Germany). Column for purification were purchased by Cytiva Life Science (Upsalla, Sweden). In general, all reagents were purchased by Sigma Aldrich (Steinheim, Germany) in at least biochemical grade. Animal experiments were performed by Oncotest GmbH (Freiburg, Germany). Amino acid analysis in serum were performed externally at MembraPure GmbH (Henningsdorf, Germany).

**Methods**

**SDS-PAGE**

Proteins and protein conjugates were analyzed by gradient tris-glycine SDS-polyacrylamide gel electrophoresis. Staining of the gel was performed by Coomassie brilliant blue G-250.

**Endotoxin amount**

Endotoxin amount of PEGylated APIT was tested at dilutions of 1:1,000, 1:10,000 and 1:50,000 using ENDOZYME® II Biomerieux Endotoxin Detection Assay, recommended spike test was performed. In the samples of dilutions of 1:10,000 and 1:50,000 of APIT-PEG no endotoxin amount could be found over the detection limit. Endotoxin units were therefore calculated in triplets of the 1:1,000 dilution of APIT-PEG and resulted in a concentration of 9 EU/mg, 0.14 EU/U. Masking of endotoxin by formulated APIT-PEG was controlled by spiking probes with endotoxins, recovery rate was 103-110%, concluding formulated APIT-PEG is not masking endotoxins.

**Animal health and monitoring**

All experiments were conducted according to the guidelines of the German Animal Welfare Act. Animal health was examined prior to tumor implantation and randomization to ensure that only animals with unobjectionable health were selected to enter testing procedures. Mice were allowed an acclimatization period of at least seven days after their arrival and prior to start of any experiment. During the experiments, animals were monitored regarding tumor burden, general condition, feed and water supply.

**Tumor sample**

The solid head and neck patient tumor HNXF 536 was derived from surgical specimens from patients at the University Hospital Freiburg (Germany) by direct implantation (passage 1), they were passaged until establishment of stable growth patterns. Master stocks of early passage xenografts were frozen in liquid nitrogen.

**Tumor volume**

The absolute volumes of subcutaneously growing tumor were determined by two-dimensional measurements with a digital caliper on the day of randomization and then twice a week. Tumor volumes were calculated according to the formula (l x w²) x 0.5, where l represents the largest diameter and w the perpendicular diameter.

**Body weights**

Animals were weighed daily during tolerability tests for one week. During efficacy tests and after the end of treatments animals were weight twice a week. If the body weight loss exceeded 10% animals were weight daily and access to food was facilitated.

**Diet and water supply**

The animals were fed autoclaved Teklab Global Extruded 19% Protein Rodent Diet (Envigo RMS SARL, Gannat, France) and had access to sterile filtered and acidified tap water pH 2.5, which was changed twive weekly. Feed and water were provided *ad libitum*. When necessary, a nutrient fortified water gel (DietGel Recovery from ClearH_2_O, Maine, USA) was provided to animal’s cages and changed at least every other day.

**DNA and Protein sequence of APIT**

```
    atg gac ggt gtc agc aga aac aga cgt caa tgt aac aga gag gtg tgc ggt tct acc tac
1   M   D   G   V   S   R   N   R   R   Q   C   N   R   E   V   C   G   S   T   Y
    gat gtg gcc gtc gtg ggg gcg ggg cct ggg gga gct aac tcc gcc tac atg ctg agg gac
20  D   V   A   V   V   G   A   G   P   G   G   A   N   S   A   Y   M   L   R   D
    tcc ggc ctg gac atc gct gtg ttc gag tac tca gac cga gtg ggc ggc cgg ctg ttc acc
40  S   G   L   D   I   A   V   F   E   Y   S   D   R   V   G   G   R   L   F   T
    tac cag ctg ccc aac aca ccc gac gtt aac ctg gag att gga ggc atg agg ttc atc gaa
60  Y   Q   L   P   N   T   P   D   V   N   L   E   I   G   G   M   R   F   I   E
    ggc gcc atg cac agg ctc tgg agg gtc att tca gaa ctc ggc cta acc ccc aag gtg ttc
80  G   A   M   H   R   L   W   R   V   I   S   E   L   G   L   T   P   K   V   F
    aag gaa ggt ttc ggc aag gag ggc aga caa aga ttc tac ctg cgg gga cag agc ctg acc
100 K   E   G   F   G   K   E   G   R   Q   R   F   Y   L   R   G   Q   S   L   T
    aag aaa cag gtc aag agt ggg gac gta ccc tat gac ctc agc ccg gag gag aaa gaa aac
120 K   K   Q   V   K   S   G   D   V   P   Y   D   L   S   P   E   E   K   E   N
    cag gga aat ctg gtc gaa tac tac ctg gag aaa ctg aca ggt cta caa ctc aac ggc gag
140 Q   G   N   L   V   E   Y   Y   L   E   K   L   T   G   L   Q   L   N   G   E
    ccg ctc aaa cgt gag gtt gcg ctt aaa cta acc gtg ccg gac ggc aga ttc ctc tat gac
160 P   L   K   R   E   V   A   L   K   L   T   V   P   D   G   R   F   L   Y   D
    ctc tcg ttt gac gaa gcc atg gat ctg gtt gcc tcc cct gag ggc aaa gag ttc acc cga
180 L   S   F   D   E   A   M   D   L   V   A   S   P   E   G   K   E   F   T   R
    gac acg cac gtc ttc aca gga gag gtc acc ctg gac gcg tcg gct gtc tcc ctc ttc gac
200 D   T   H   V   F   T   G   E   V   T   L   D   A   S   A   V   S   L   F   D
    gac cac ctg gga gag gac tac tat ggc agt gag atc tac acc cta aag gaa gga ctg tct
220 D   H   L   G   E   D   Y   Y   G   S   E   I   Y   T   L   K   E   G   L   S
    tcc gtc cca caa ggg ctc cta cag gct ttt ctg gac gcc gca gac tcc aac gag ttc tat
240 S   V   P   Q   G   L   L   Q   A   F   L   D   A   A   D   S   N   E   F   Y
    ccc aac agc cac ctg aag gcc ctg aga cgt aag acc aac ggt cag tat gtt ctt tac ttt
260 P   N   S   H   L   K   A   L   R   R   K   T   N   G   Q   Y   V   L   Y   F
    gag ccc acc acc tcc aag gat gga caa acc aca atc aac tat ctg gaa ccc ctg cag gtt
280 E   P   T   T   S   K   D   G   Q   T   T   I   N   Y   L   E   P   L   Q   V
    gtg tgt gca cag aga gtc atc ctg gcc atg ccg gta tac gct ctg aac caa cta gac tgg
300 V   C   A   Q   R   V   I   L   A   M   P   V   Y   A   L   N   Q   L   D   W
    aat cag ctc aga aat gac cga gcc acc caa gcg tac gct gcc gtt cgc ccg att cct gca
320 N   Q   L   R   N   D   R   A   T   Q   A   Y   A   A   V   R   P   I   P   A
    agt aag gtg ttc atg acc ttt gat cag ccc tgg tgg ttg gag aac gag agg aaa tcc tgg
340 S   K   V   F   M   T   F   D   Q   P   W   W   L   E   N   E   R   K   S   W
    gtc acc aag tcg gac gcg ctt ttc agc caa atg tac gac tgg cag aag tct gag gcg tcc
360 V   T   K   S   D   A   L   F   S   Q   M   Y   D   W   Q   K   S   E   A   S
    gga gac tac atc ctg atc gcc agc tac gcc gac ggc ctc aaa gcc cag tac ctg cgg gag
380 G   D   Y   I   L   I   A   S   Y   A   D   G   L   K   A   Q   Y   L   R   E
    ctg aag aat cag gga gag gac atc cca ggc tct gac cca ggc tac aac cag gtc acc gaa
400 L   K   N   Q   G   E   D   I   P   G   S   D   P   G   Y   N   Q   V   T   E
    ccc ctc aag gac acc att ctt gac cac ctc act gag gct tat ggc gtg gaa cga gac tcg
420 P   L   K   D   T   I   L   D   H   L   T   E   A   Y   G   V   E   R   D   S
    atc ccg gaa ccc gtg acc gcc gct tcc cag ttc tgg aca gac tac ccg ttt ggc tgt gga
440 I   P   E   P   V   T   A   A   S   Q   F   W   T   D   Y   P   F   G   C   G
    tgg atc acc tgg agg gcc ggc ttc cat ttc gat gac gtc atc agc acc atg cgt cgc ccg
460 W   I   T   W   R   A   G   F   H   F   D   D   V   I   S   T   M   R   R   P
    tca ctg aaa gat gag gta tac gtg gtg gga gcc gac tac tcc tgg gga ctt atc tcc tcc
480 S   L   K   D   E   V   Y   V   V   G   A   D   Y   S   W   G   L   I   S   S
    tgg ata gag ggc gct ctt gag acc tcg gaa aac gtc atc aac gac tac ttc ctc tga
500 W   I   E   G   A   L   E   T   S   E   N   V   I   N   D   Y   F   L   *
```

Table SI 1: Design and dosing of efficacy experiment

| Group ID | Compound | Total daily dose | Dosing days | No. of animals |
| --- | --- | --- | --- | --- |
| 1 | Control Vehicle | 5 ml/kg | 1, 3, 5, 7, 9, 11, 13, 15, 17, 19, 21 | 10 |
| 2 | Doxorubicin | 6 mg/kg | 1, 8, 15, 22 | 10 |
| 3 | APIT-PEG | 15 U/kg | 1, 3, 5, 7, 9, 11, 13, 15, 17, 19, 21 | 8 |
| 4 | APIT-PEG | 40 U/kg | 1, 3, 5, 7, 9, 11, 13, 15, 17, 19, 21 | 6 |

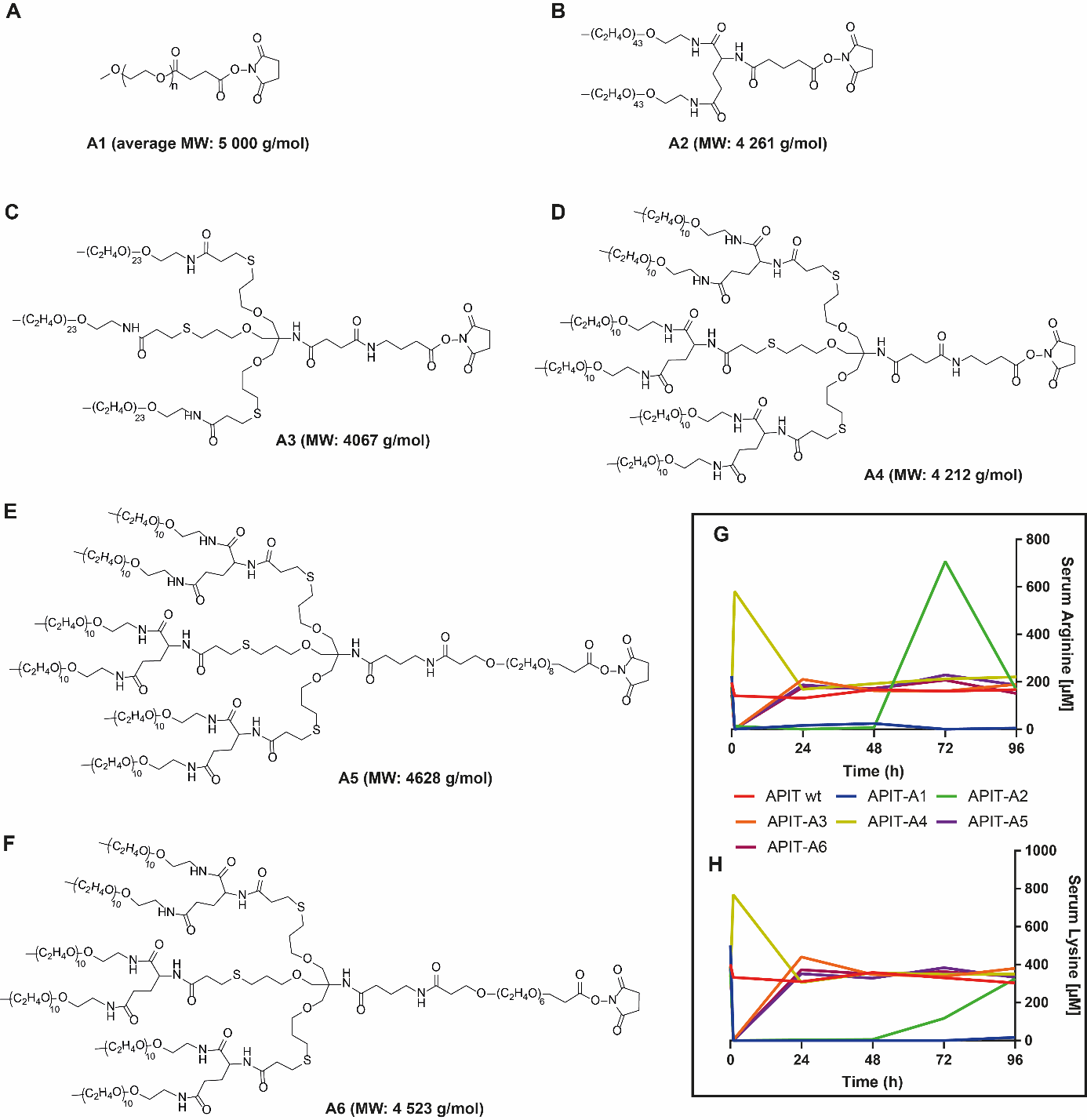

Figure S1: Screening of APIT-PEG conjugates on the pharmacodynamic. Molecular structure of A) polydisperse linear PEG-NHS with an average molecular mass of 5 000 g/mol, B) polydisperse 2-Arm PEG-NHS with a molecular weight of 4 261 g/mol, C) mono-disperse 3-Arm PEG-NHS with a molecular weight of 4 067 g/mol, D) mono-disperse 6-Arm PEG NHS with a molecular weight of 4 212 g/mol, E) mono-disperse 6-Arm PEG-Linker(8) NHS with a molecular weight of 4 628 g/mol, F) mono-disperse 6-Arm PEG-Linker(6) NHS with a molecular weight of 4 523 g/mol, and serum analysis after a single administration of APIT PEG-conjugates with A1-A6 of after 1 h, 24 h, 48 h, 96 h of G) arginine level and H) lysine level showing a throughout depletion of arginine and lysine for up to 96 h with APIT-A1 (APIT-PEG5kDa) and a depletion for up to 48 h of APIT-A2 (APIT‑2‑Arm‑PEG4.3kDa), whereas the other PEG conjugates did show initial depletion but serum concentrations were restored within 24 h.

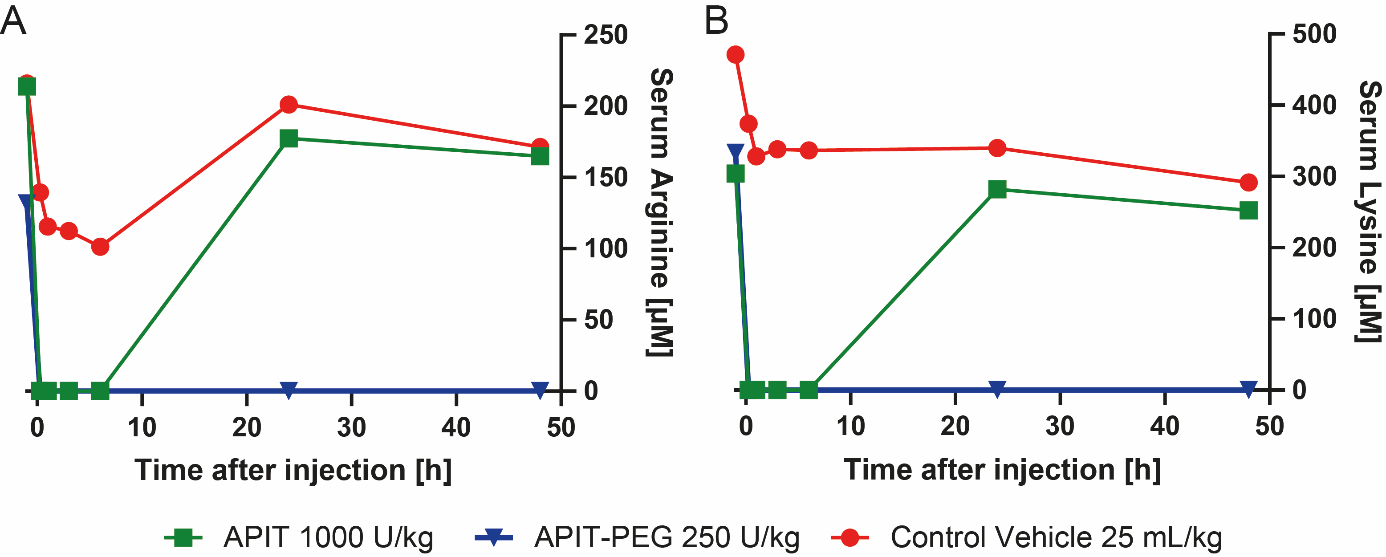

Figure S2: Pharmacodynamics of a single administration of 1000 U/kg APIT or 250 U/kg APIT-PEG on A) arginine concentration in serum and B) lysine concentration in serum showing a prolonged serum depletion of 1000 U/kg APIT compared to 250 U/kg for at least 3h, amino acid recovery was observed after 24h after the injection of 1000 U/kg APIT.

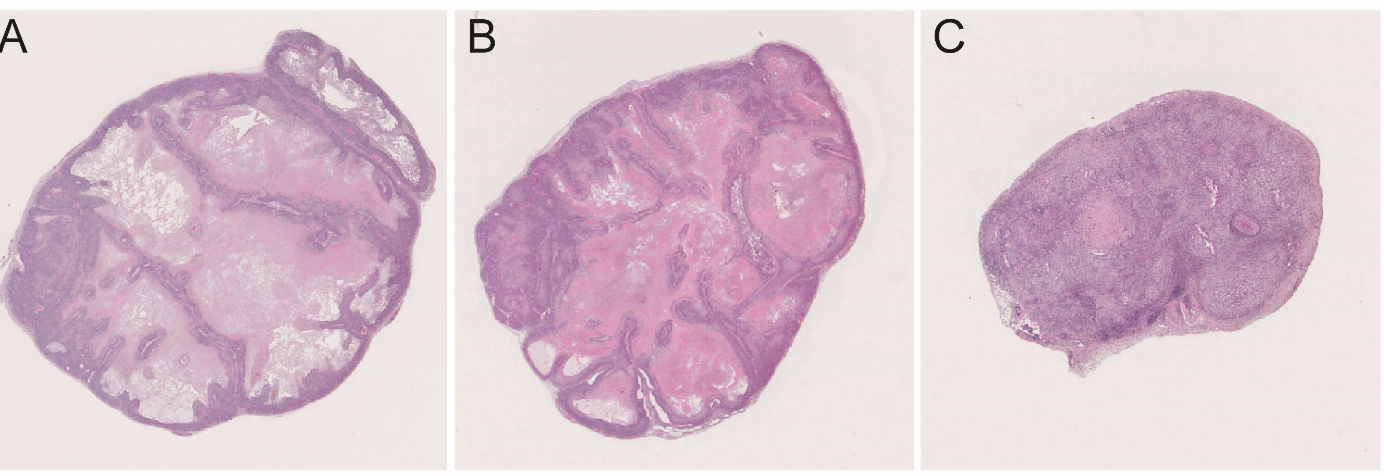

Figure S3: Representative histology (H&E staining) of patient derived head and neck cancer HNXF 536 as xenograft in mice 42 days after randomization. A) of the non-treated control group B) of the group treated with 40 U/kg APIT-PEG and C) tumor tissue of the 40 U/kg APIT-PEG treated mouse showing a regression of tumor.

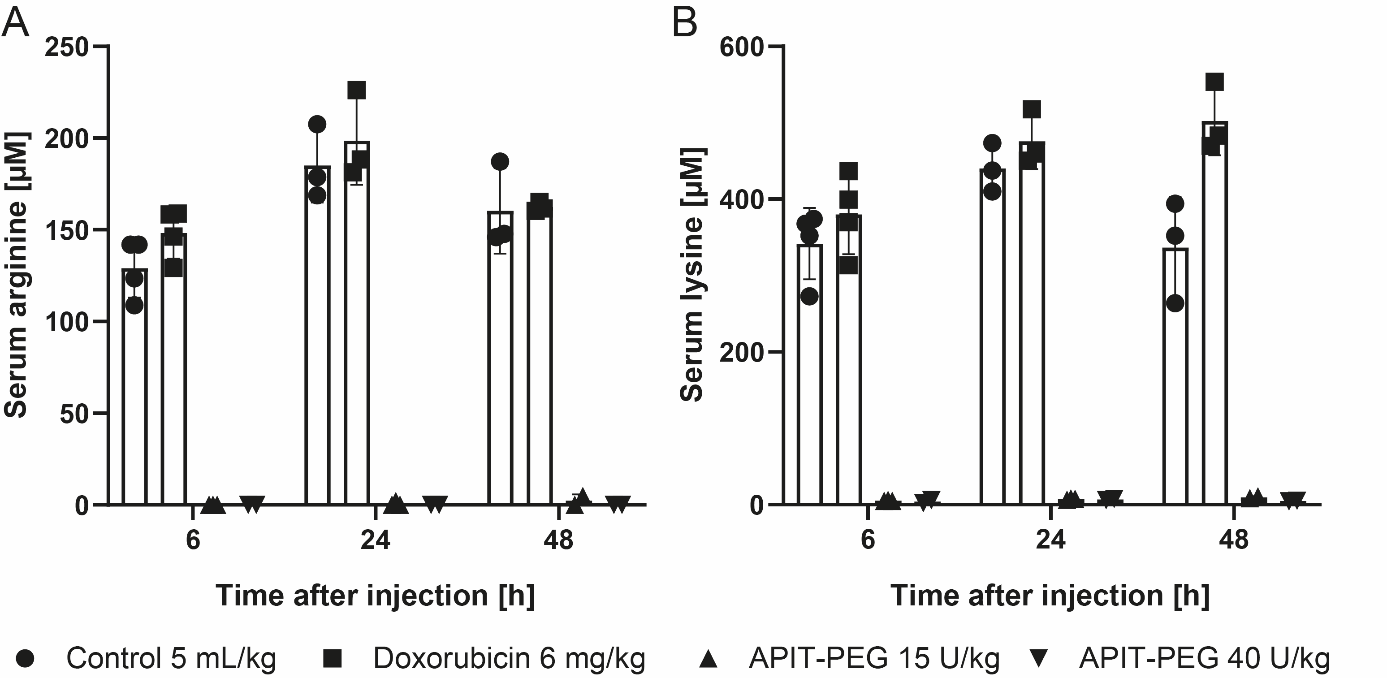

Figure S4: Mean and SD of amino acid concentration in serum of treated xenografted HNX 536-mice A) Concentration of serum arginine and B) Concentration of serum lysine showing a complete depletion of arginine and lysine of APIT-PEG treated mice. However, an beginning increase of arginine amount to 4.8 µM in one mouse of the APIT-PEG 15 U/kg was detectable.

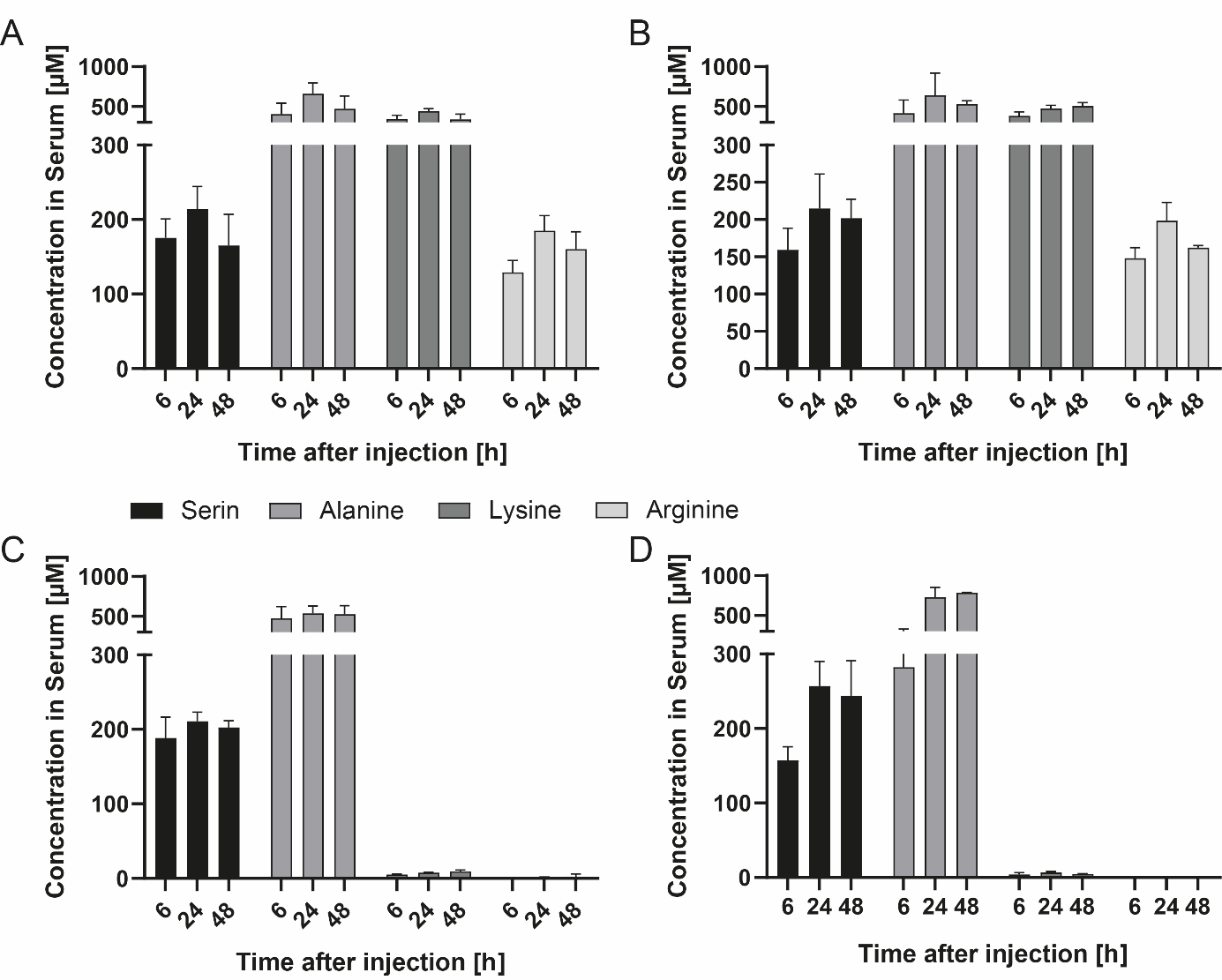

Figure S5: Mean and SD of serin, alanine, lysine and arginine concentration in serum of xenografted HNX 536-mice of the A) Non-treated control group B) Doxorubicine treated group C) APIT-PEG 15 U/kg and D) APIT-Peg 40 U/kg group. Showing a depletion of arginine and lysine in the APIT-PEG treated group, whereas Serin and Alanine levels remained unchanged. An increase arginine level of 4.8 µM was seen in one 48 h sample of APIT-PEG 15 U/kg mouse.

An increase arginine level of 4.8 µM was seen in one 48 h sample of APIT-PEG 15 U/kg mouse.

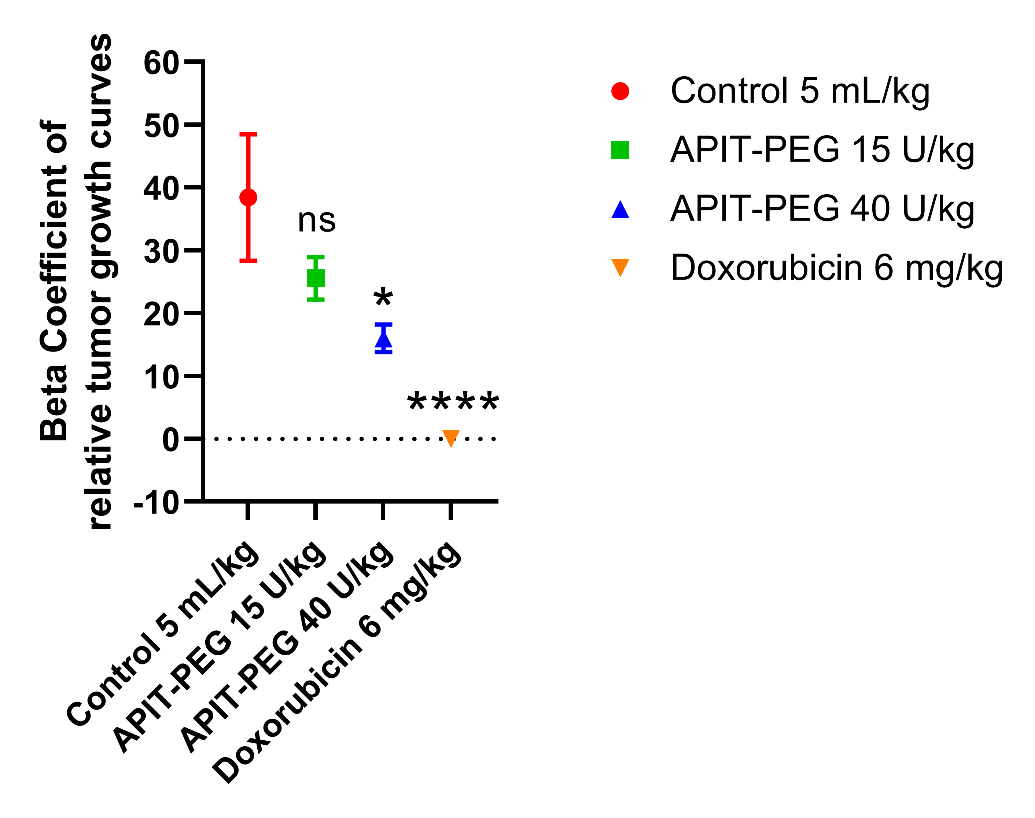

Figure S6: Comparison of beta coefficients of the curves of the relative tumor growth performed by a linear regression fit. The beta coefficients are shown as mean with the standard deviation. Statistical analysis was performed by ordinary One-way ANOVA in a multiple comparison to the control group. The curve of the relative tumor growth of 15 U/kg was not significant, whereas the 40 U/kg (p = 0.0493) and doxorubicin (p < 0.0001) treatment curves were significantly different to the control group
